# BSImp: imputing partially observed methylation patterns for evaluating methylation heterogeneity

**DOI:** 10.1101/2021.12.07.471020

**Authors:** Ya-Ting Chang, Ming-Ren Yen, Pao-Yang Chen

## Abstract

DNA methylation is one of the most studied epigenetic modifications that has applications ranging from transcriptional regulation to aging, and can be assessed by bisulfite sequencing (BS-seq) at single base-pair resolution. The permutations of methylation statuses at bisulfite converted reads reflect the methylation patterns of individual cells. These patterns at specific genomic locations are sought to be indicative of cellular heterogeneity within a cellular population, which are predictive of developments and diseases; therefore, methylation heterogeneity has potentials in early detection of these changes. Computational methods have been developed to assess methylation heterogeneity using methylation patterns formed by four CpGs, but the nature of shotgun sequencing often give partially observed patterns, which makes very limited data available for downstream analysis. While many programs are developed to impute methylation levels genomewide, currently there is only one method developed for recovering partially observed methylation patterns; however, the program needs lots of data to train and cannot be used directly; therefore, we developed a probabilistic-based imputation method that uses information from neighbouring sites to recover partially observed methylation patterns speedily. It is demonstrated to allow for the evaluation of methylation heterogeneity at three times more regions genome-wide with high accuracy for data with moderate depth. To make it more user-friendly we also provide a computational pipeline for genome-screening, which can be used in both evaluating methylation levels and profiling methylation patterns genomewide for all cytosine contexts, which is the first of its kind. Our method allows for accurate estimation of methylation levels and makes evaluating methylation heterogeneity available for much more data with reasonable coverage, which has important implications in using methylation heterogeneity for monitoring changes within the cellular populations that were impossible to detect for the assessment of development and diseases.

## 1 INTRODUCTION

Methylation is one of the most studied epigenetic modifications (Moore et al., 2013). It is known to be involved in a wide range of key biological processes including regulation of gene expression, developments (Hsieh et al., 2020), aging and silencing of transposable elements (Jin et al., 2011). The study of methylation at single nucleotide resolution is made possible through next generation sequencing when it is coupled with bisulfite treatment (Barros-Silva et al., 2018). Methylation level is used extensively in comparing between samples of different conditions (Hsieh et al., 2020) and their correlation with gene expression is usually studied (Hanley et al., 2017). When looking at reads covering multiple cytosines there are also methylation patterns, or permutations of methylation statuses spanning multiple cytosines in a row. As one read represents a cell within a bulk sequencing data, it is sought that methylation patterns can be used to study cellular heterogeneity, which was discovered to be associated with diseases such as cancer (Jin et al., 2011); the more heterogeneous the tumours are, the worse the clinical outcomes (Landau et al., 2014). A few methods had been proposed to study cellular heterogeneity including experimentally using single cell bisulfite sequencing (scBS-seq); however, there are significant challenges associated in that it is technical difficult to isolate individual cells let alone bisulfite treatment can also destroy the DNA sample. A new method is to quantify methylation heterogeneity using methylation patterns that are formed by methylation statuses of several cytosines within the same reads in bulk sequencing data (Zhang et al., 2011; Shannon, 1948; Hill, 1973). However, owing to the nature of shotgun sequencing and average depth and coverage of data given by most whole genome bisulfite sequencing data (WGBS) and reduced representation bisulfite sequencing (RRBS) data due to financial consideration, it is still challenging to measure methylation heterogeneity accurately as even depths do not constitute enough methylation patterns at each cytosine.

Imputation is a commonly used technique to overcome this type of problems; However, most imputation methods developed for methylation data analysis such as METHimpute (Taudt et al., 2018), Melissa (Kapourani and Sanguinetti, 2019) and DeepCpG (Angermueller et al., 2017) were developed for methylation levels of individual CpG sites or within specific regions (promoters) for single cell methylomes where data are sparse (DeepCpG, Melissa) or WGBS (METHimpute). Despite their usefulness in inferring methylation levels genomewide, they were not designed for and hence are unable to recover read specific methylation patterns that are needed for the estimation of methylation heterogeneity since it requires read identity for each methylation status. PReLIM (Scott et al., 2020), on the other hand, attempts to impute methylation statuses on individual sequencing reads; however, the method requires training models using many bins and the program written is not straightforward.

DNA methylation is catalysed by a family of DNA methyltransferases (DNMTs) (Jin et al., 2011). Different contexts of methylation, methylation occurring at CG, CHG, and CHH contexts where H is any of A,C, or T, are responsible by different groups of DNMTs. DNA methylation for mammalians primarily occur at CG (Jin et al., 2011) while methylation at other contexts CHG and CHH are also common for plants although their roles are not clear. Currently, most of the studies based on methylation heterogeneity are for human diseases, hence, only available for CG methylation but the same concept can be useful for other contexts as well, for example, for studying DNMTs and the pathway involved (Harris and Zemach, 2020). Therefore, it is our aim to develop an imputation method that is accurate, speedy and produces outputs widely applicable to animals as well as plants and fungi, with higher resolution (methylation pattern information).

There is high correlation of methylation among cytosines that are nearby (Affinito et al., 2020). We use this property extensively to borrow the most information from nearby sites and developed a probabilistic-based imputation method to impute accurate methylation statuses speedily. Our program is the first of its kind to be able to take any methylation contexts (not limited to CG) that has accuracy comparable with the only existing method that imputes methylation statuses. It also has the flexibility for user to specify window size in number of cytosines fixed across the genome for imputation and genomewide profiling. After all, it is easier to use, can be run with one command and outputs results readily for downstream analyses.

## 2 METHOD

Considering methylation patterns formed by methylation statuses of multiple successive cytosines of the same reads, there is usually missing values of methylation statuses within a window of fixed size (number of cytosines). We assume that the pattern of methylation is similar for cells within a population and that the behaviour of cells or the methylation statuses of a cell at a given position can be predicted by those statuses nearby and cells nearby; therefore, using law of total probability, let the methylation status of a cytosine at a position *j* for read *i* be *m*_*ij*_, then the probability of *m*_*ij*_ being methylated, or 1, is

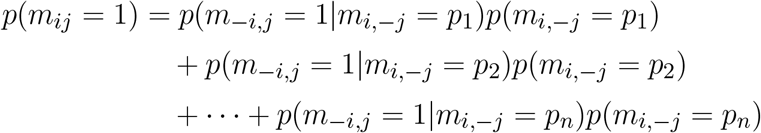

where *p*_*i*_ are subpatterns of complete patterns within the same window, or methylation patterns at positions other than *j* and *p*(*m*_*−i,j*_ = 1|*m*_*i,−j*_ = *p*_*s*_) is the observed probability of cytosines being methylated at position *j* given subpattern *m*_*i,−j*_ within the window is like *p*_*s*_. The reads eligible for imputation is specified to be those missing at most 1 methylation status within the window. Since *m*_*ij*_ is the only missing value in the window for the same read, *m*_*i,−j*_ must equal to one of *p*(*m*_*−i,j*_ = 1|*m*_*i,−j*_ = *p*_*s*_)*p*(*m*_*i,−j*_ = *p*_*s*_) where *s* ⊂ {1, 2, …, *n*}. However, if the subpattern is not observed, or there is no complete pattern with subpattern that resembles *m*_*i,−j*_, it is taken as the methylation level at position *j*, or *p*(*m*_*tj*_ = 1) for all reads *t* that are observed at position *j*. An illustration of the eligibility of reads for imputation and a possible imputation result can be found in Figure 1.

**Figure 1.**
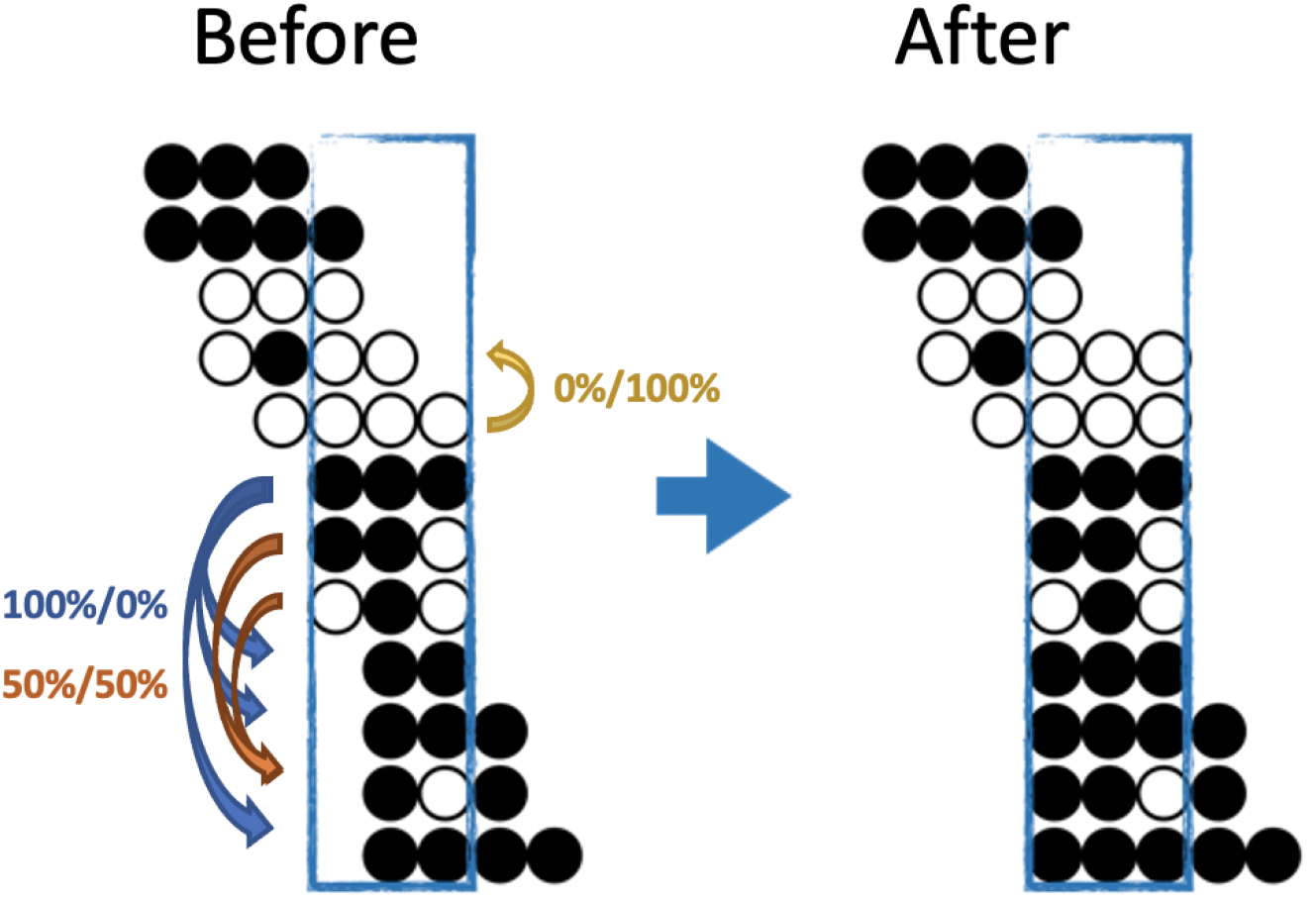
An illustration of both the eligibility and a possible imputation result for BSImp. Given we are interested in window size of three cytosines, a region is selected as enclosed by a blue rectangle. Each line of dots represents a read aligned to a specific genomic region; black (white) dots represents a methylated (unmethylated) cytosine for the given read. Four complete patterns are observed, which makes the window eligible for imputation. Within the window, only reads missing at most one methylation statuses (dots) are eligible for imputation; there are five in the example. Looking at the topmost reads with one missing pattern, the rest of the pattern resembles that of the complete pattern below, so it has a probability of zero being methylated. The second last read has methylation pattern (black, white in the second and third position) resembles two other reads, which have methylation statuses of methylated and unmethylated, one each, so it has 50% of being either.

In our implementation, the imputations are done alongside genome screening where windows of fixed size of cytosines of the same methylation contexts are extracted, imputed if valid and profiled for their copy numbers of methylated, unmethylated reads and every possible methylation patterns. It is done through sliding windows with one cytosine overlapping. Only windows with at least two complete patterns are considered for imputation and results outputted if a given cytosine has enough depths as specified by the user.

## 3 RESULT

To evaluate imputation performances, different types of data including whole genome bisulfite sequencing (WGBS) and reduced representation bisulfite sequencing (RRBS) data are used to assess the effect of imputation on genome coverage and the accuracy of prediction evaluated using both methylation statuses and methylation level. The WGBS data selected are from Arabidopsis thaliana and RRBS data is from human. In the evaluation only the forward strand is used.

### 3.1 Imputation can increase significantly in coverage

The primary purpose of imputation is to increase coverage genomewide for downstream analysis so we first examine the increase in coverage of our method. Two WGBS datasets with average depths of 18x are used. Data of lower depths are obtained by cutting off the head and tail of each read to reach desirable average depths. The chosen depths are 5x, 8x, 10x, 15x and 18x. We can see a clear trend of increase in coverage as depth increases and the coverage for imputed methylomes are much higher than that before imputation with maximum linear increase between 18 and 20% for depths between 8x and 10x. Looking at Figure 2A and Figure 2B we also see the coverage for methylation level is much higher than methylation heterogeneity; there are two reasons for this: the depths required for evaluating methylation level is 4 and for evaluating methylation heterogeneity is 8 and it is much harder to observe complete methylation patterns compared to reads at individual cytosines; hence, the need for imputation for the evaluation of methylation heterogeneity.

**Figure 2.**
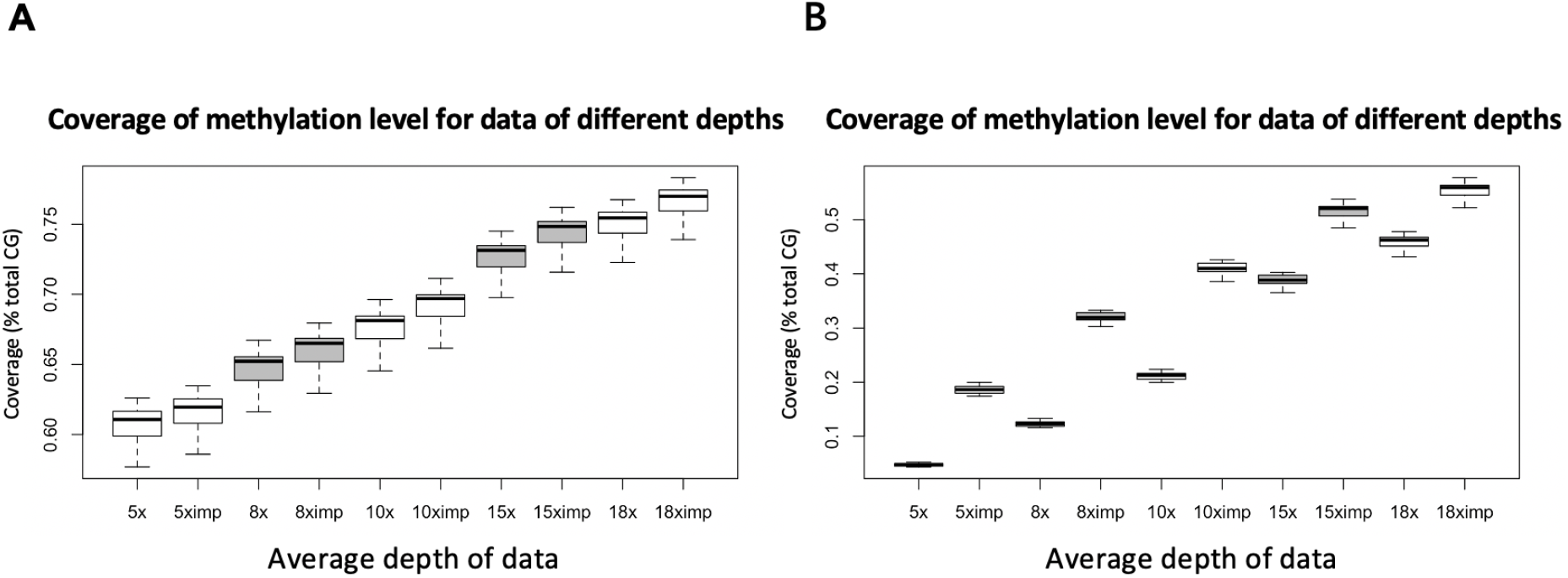
An illustration of increases in coverage for the evaluation of methylation level and methylation heterogeneity for WGBS data with common depths. A) Coverage (% total CG) of possible evaluation of methylation level before and after imputation evaluated with requirement of minimum depth of 4 reads at each cytosine. B) Coverage (% total CG approximately) of possible evaluation of methylation heterogeneity before and after imputation using window size of 3 CpG and minimum 8 reads per window. Eight was chosen for it is the minimum number of reads required for us to see all possible patterns, if they all appear. Each boxplot is based on 5 chromosomes and 2 libraries (WGBS) (10 data points) so % total CG is the number of the CG sites in the chromosomes. Only the forward strand is considered. Since methylation patterns are profiled using sliding windows of one cytosines, the total number of windows we can get is slightly fewer than the total number of CG sites.

### 3.2 Imputation predicts methylation statuses accurately

Given imputation increases coverage, it is also important to know how much bias it introduces. We first compare our method with PReLIM which is the only existing method that also recovers methylation patterns. The result as shown in Figure 3A is obtained by getting all complete patterns within windows of 4 CpGs in chromosome 2 of a human cancer data, removing multiple methylation statuses at random within each window and impute the missing values using different methods (BSImp, PReLIM and column mean) for comparison. Column mean is a method that uses methylation level for the genomic position as the probability of methylation for all reads eligible for imputation. The accuracy is calculated as the mean number of correctly imputed methylation statuses. Figure 3A shows that our method has higher accuracy (over 85%) than PReLIM and using column mean as probability for predicting methylation statuses.

**Figure 3.**
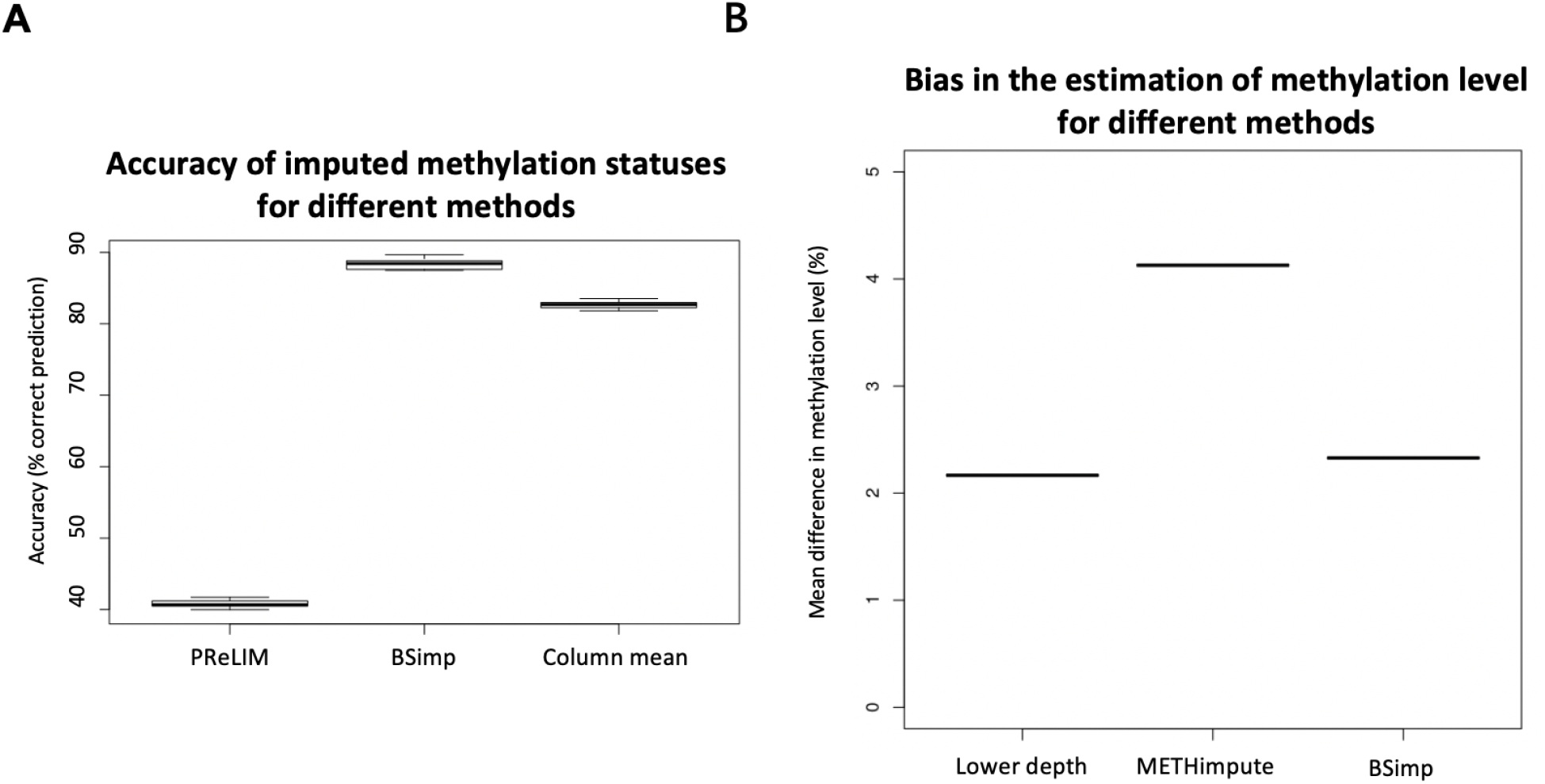
Evaluation of accuracy of BSimp by comparing with existing methods. (A) Boxplot of accuracy of methylation statuses imputed compared with PReLIM and column mean. Non-overlapping windows of four CGs are extracted, multiple methylation statuses removed at random and imputed using all three methods to calculate mean accuracy of prediction for each window. PReLIM needs training of models, (complete parts of) all windows included in the evaluation were used for training. Column mean is a method that uses methylation level for the genomic position as the probability of methylation for all reads eligible for imputation. The results are based on 783 windows. (B) Boxplot of mean difference in methylation level of common regions compared with METHimpute and data of lower depths. Four libraries were obtained by downsampling 50% of reads independently and imputed using METHimpute and BSImp. Methylation levels obtained before and after imputation are compared with library before downsampling as treating it as the correct answer. Only common regions (CG) are considered. Each boxplot is based on four results.

Since imputation changes the estimate for methylation level at each cytosine, we also assess the accuracy (bias) using absolute changes in methylation level estimated. Figure 3B is obtained by calculating the mean absolute difference in methylation level across common regions with estimate of methylation level between original data (18x) and data of lower depth by downsampling 50% of reads and imputed data using different methods as indicated by the x-axis. Four libraries are obtained by downsampling 50% of the reads; each boxplot consists of result from 4 libraries. Figure 3B shows our method does not create much difference in methylation level compared to data before imputation as indicated by lower depth and METHimpute introduces much larger bias for these cytosines.

## 4 DISCUSSION

There is only one existing method that recovers methylation patterns, which can be beneficial for the evaluation of methylation heterogeneity; however, the program written is standalone; it only imputes or completes a binary matrix of indicator variables that represent the methylation statuses within a window of given numbers of CpGs; it is up to the users to extract the windows for training and predicting and to output results useful for downstream analyses. On the other hand, our program (Figure 4) is able to screen for methylation pattern genomewide, impute missing statuses and output the profiles of methylation statuses at each cytosine and the copy number of every possible methylation patterns given the size of the window. In other words, it is the first of its kind and all in one. The results produced include number of methylated and unmethylation cytosine at each position given the depth is enough and the copy numbers of every possible methylation patterns starting at the same position, which can be easily used to evaluate methylation levels and methylation heterogeneity. We compared the accuracy of BSImp in terms of accurate prediction of methylation statuses with PReLIM as it is the only method that recovers methylation patterns and to our surprise, PReLIM performs a lot worse and we also ran an analysis of the breakdown of methylation levels of the windows we used in the evaluation and it turns out that PReLIM performs bad when methylation levels are high, which might requires some tuning of parameters as high methylation level can be common for most methylation contexts of interest; i.e., CpG for human.

**Figure 4.**
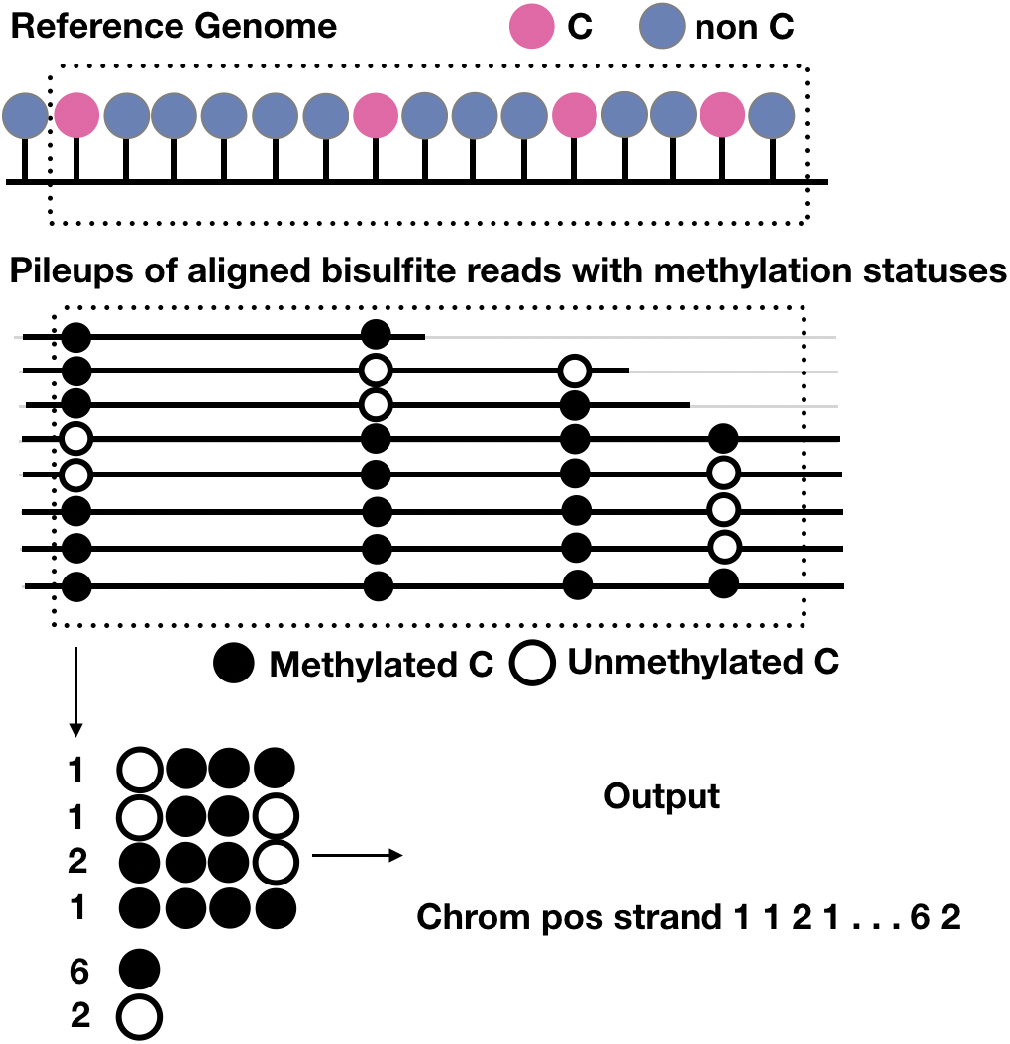
Schematic representation of the output of BSImp. First the reads are aligned to the reference genome, given any window of specified number of cytosines (here is 4), methylation statuses are extracted and missing values imputed if valid, then methylation profiled as a vector including chromosome, position, strand, the copy numbers of all possible methylation patterns starting at the position, and number of methylated and unmethylated cytosines.

As for methylation, existing methods that impute methylation levels were mostly developed for imputing whole methylome of sparse data such as single cell methylomes; however, METHimpute is developed for imputing methylation level of entire methylomes using WGBS, which is closer to our aim; therefore, we only compared with METHimpute using the sites with common coverage (of METHimpute and BSImp using WGBS) where BSImp has lower and the result indicates our method is comparable to METHimpute at predicting methylation level and the bias is only slightly larger than that of lower depth (data before imputation) by treating original data (not downsampled) as target.

## 5 ADDITIONAL REQUIREMENTS

### AUTHOR CONTRIBUTIONS

Y.C. conceived the study, collected the data, and wrote the manuscript. P.C. supervised the work, revised the manuscript and provided the funding for this work. M.Y. supervised the development of the programs, was in charge of the evaluation and maintains the repository.

### FUNDING

This work was supported by grants from Academia Sinica, and Ministry of Science and Technology Taiwan (109-2313-B-001-009-MY3 and 108-2313-B-001-013-MY3) to P.-Y. C.

## ACKNOWLEDGMENTS

Not applicable.

## DATA AVAILABILITY STATEMENT

- All sequencing data used in the study were downloaded from the NCBI Gene Expression Omnibus under accession numbers GSE81407 for Arabidopsis Hsieh et al. (2020) and GSE95656 for colorectal cancer (CRC) Hanley et al. (2017) and are publicly available as of the date of publication. All other data reported in this paper will be shared by the lead contact upon request.
- All original code has been deposited at https://github.com/britishcoffee/BSimp and is publicly available as of the date of publication..
- Any additional information and code required to reproduce the results reported in this paper is available from the lead contact upon request.

